# First report of pathogenic *Leptospira* in Sri Lankan bats: A potential reservoir risk

**DOI:** 10.1101/2024.09.27.615370

**Authors:** Thejanee Perera, Franziska Schwarz, Therese Muzeniek, Sahan Siriwardana, Beate Becker-Ziaja, Inoka Perera, Shiroma Handunnetti, Jagathpriya Weerasena, Gayani Premawansa, Sunil Premawansa, Wipula Yapa, Claudia Kohl, Andreas Nitsche

## Abstract

**Background:** Leptospirosis is a bacterial infection caused by spirochetes of the genus *Leptospira*. It affects both humans and animals, posing a significant public health threat worldwide. In Sri Lanka, leptospirosis causes annual outbreaks, particularly during the monsoon seasons. The disease is often difficult to diagnose and can have a high mortality rate. Bats are potential reservoirs for *Leptospira* in other countries. This study investigated the presence and diversity of *Leptospira* in bats inhabiting the Wavulgalge, a natural cave in Sri Lanka.

**Methodology/Principal Findings:** Using PCR and Sanger sequencing techniques, *Leptospira borgpetersenii* and *Leptospira kirschneri* were identified in *Miniopterus fuliginosus* and *Rousettus leschenaultii* bats. Out of 148 bat urine samples, 16 tested positive via RT-qPCR, with *M. fuliginosus* samples harboring *L. borgpetersenii* and one *R. leschenaultii* sample harboring *L. kirschneri* revealed by Sanger sequencing method.

**Conclusions/Significance:** These findings enhance the understanding of the zoonotic role of Sri Lankan bats in *Leptospira* transmission.

**Author Summary:** Leptospirosis is a bacterial disease that affects both humans and animals, particularly in tropical regions. It spreads through contact with water or soil contaminated by the urine of infected animals. In Sri Lanka, leptospirosis poses a significant public health challenge, with frequent outbreaks during the rainy season. However, while much research has focused on domestic animals as potential carriers, the role of bats in the transmission of this disease has not been fully explored.

In this study, we investigated whether bats in Sri Lanka can act as carriers of *Leptospira* bacteria, which could contribute to the spread of leptospirosis to humans. We focused on a population of bats from the Wavulgalge cave and tested them for the presence of *Leptospira* using molecular methods. Our results provide the first evidence that *Leptospira* is present in Sri Lankan bats, highlighting their potential role as a reservoir for the bacteria.

These findings are important because they suggest that bats, like rodents and livestock, could contribute to the spread of leptospirosis in Sri Lanka. Understanding which animals carry the bacteria helps inform public health strategies, allowing for more targeted surveillance and control measures. By including bats in these efforts, we can better prevent future outbreaks of leptospirosis and protect at-risk communities.

This study also contributes to the broader understanding of how wildlife can influence human health, especially in tropical environments where human-animal interactions are common. Our work emphasizes the need for a One Health approach, which considers the connections between human, animal, and environmental health, to tackle neglected diseases like leptospirosis.

## Introduction

Humans contract Leptospirosis through direct contact with the urine of infected animals or contaminated environment (*1*). In addition to entering the body through the mucous membranes of the mouth, nose, and eyes, *Leptospira* bacteria can also enter through cuts or abrasions on the skin (*2*). Clinical manifestations of *Leptospira* infections in humans can range from flu-like symptoms to organ failure and even death causing around 60,000 fatalities and an average of one million infections globally each year (*1,3*).

Leptospirosis causes annual outbreaks in Sri Lanka associated with the two monsoon seasons followed by floods (*4*). It was declared a notifiable disease in 1991 (*3*). The estimated case fatality rate of leptospirosis at 7% is more than 10 times higher than the case fatality rate reported for Dengue virus in Sri Lanka (*4*). The 2008 leptospirosis outbreak in Sri Lanka, which resulted in over 7,400 suspected cases and more than 200 deaths, prompted increased focus on improving diagnosis and surveillance to better understand and manage the disease (*5*). Bats are increasingly recognized as potential reservoirs of infectious agents like *Leptospira*, with Sri Lankan bats known to carry bacteria such as *Salmonella* and various viruses (*6–10*). In Sri Lanka, previous studies have identified several *Leptospira* serovars in livestock, wild and domestic animals, such as cattle, cats, dogs, pigs, elephants, shrews, rats and mice (*11–15*). However, the specific role of bats in the transmission dynamics of leptospirosis in Sri Lanka is not fully understood (*15,16*).

Therefore, the aim of this study was to identify the presence of pathogenic *Leptospira* species in Sri Lankan bats and to identify their role as potential reservoir hosts for maintaining the sylvatic cycle. The *lipL32* gene, encoding an outer membrane lipoprotein conserved among pathogenic *Leptospira* species, was targeted for initial PCR screening of bat urine samples (*17,18*). Positive samples were further screened and sequenced using primers for the *flaB*, *secY,* and 16s rRNA (*rrs2*) genes. These genes were chosen to compare sequences with previous studies (*19–22*). The *flaB* encodes flagellin, a key component of *Leptospira* motility while *secY* gene encodes a membrane protein involved in bacterial protein transport (*23,24*). 16s rRNA gene (*rrs*2) sequences are stored in GenBank for all type strains of the species of the family *Leptospiraceae*. By using these genes as molecular markers, the study aimed to establish sequence similarities and provide valuable insights into the genetic characteristics of *Leptospira* strains identified in Sri Lankan bats.

## Materials and Methods

### Sampling location

The study was conducted at Wavulgalge cave, a natural cave located in Koslanda, Monaragala district, Sri Lanka (6° 43’ 37.1676’’ N, 81° 3’ 11.7216’’ E) as shown in Fig 1.

**Fig 1.**
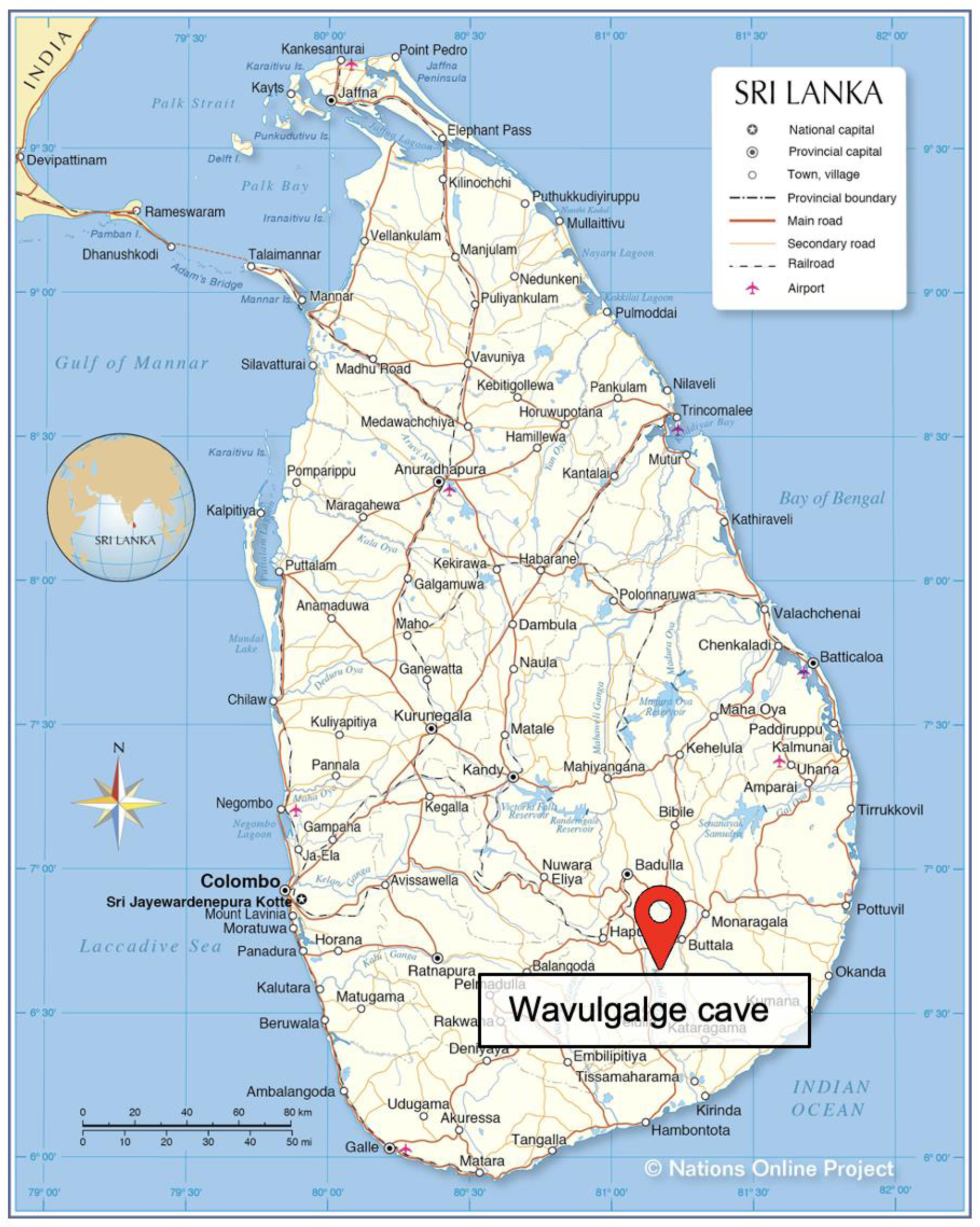
Sampling Location: Wavulgalge cave, Sri Lanka.

The location pin of the figure indicates the Wavulgalge cave, situated in Sri Lanka. Image used for educational purposes, courtesy of Nations Online Project.

This site was selected due to its significance as one of Sri Lanka’s largest sympatric bat roosts, housing diverse bat species coexisting in a sympatric colony. Fig 2 shows the interior of the Wavulgalge cave.

**Fig 2.**
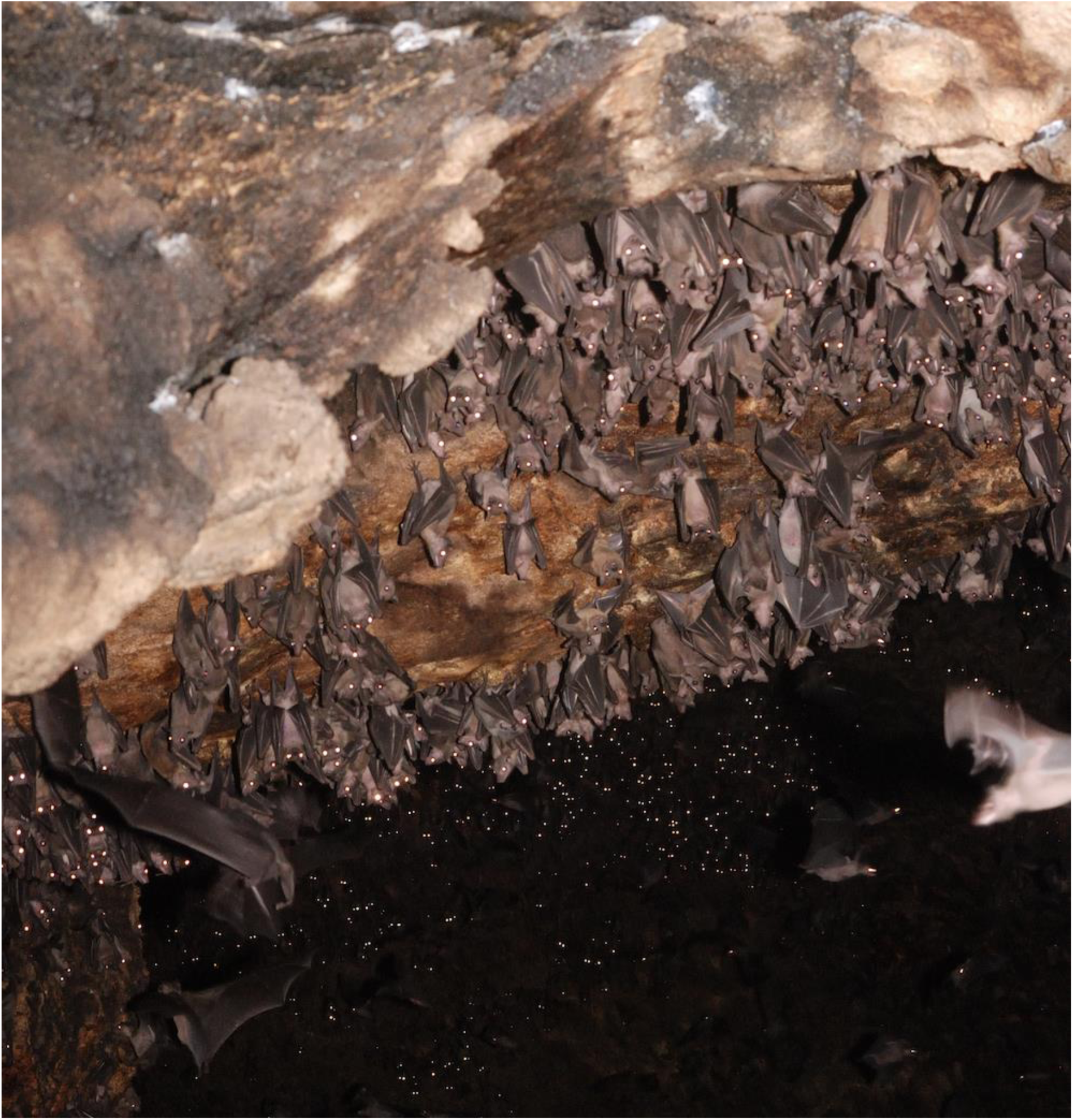
Interior view of the Wavulgalge cave, Sri Lanka. Copyright © Timothy Hornby

### Sampling period

Bats were sampled in March and July 2018, and January 2019, covering reproductive and seasonal variations. Bats were captured with hand nets during their foraging emergence, then documented for age, gender, and forearm length. Age groups were categorized into juvenile, sub-adult, and adult, with sexual dimorphism used to identify gender.

### Biological sample collection

Urine samples were collected with CleanFoam® swabs, from four bat species: *Miniopterus fuliginosus*, *Hipposideros speoris*, *Rousettus leschenaultii*, and *Rhinolophus rouxii*. During the March 2018 field visit, 11 urine samples were collected from *H. speoris*, *R. leschenaultii*, and *R. rouxii*. However, no *M.fuliginosus* bats were captured due to the low bat population. In July 2018, *M.fuliginosus* bats were more abundant due to seasonal migration, and 106 samples were collected during this visit. *H. speoris* and *R. rouxii* bats were not sampled in July, while two *R. leschenaultii* bats were sampled. The final field visit in January 2019 included all four bat species, with six urine samples collected from *H. speoris*, *R. leschenaultii*, and *R. rouxii*, along with eleven samples from *M. fuliginosus*, bringing the total to 148 urine samples across all three sampling points. Urine samples were stored in 2 ml microtubes, and transported in Va-Q-Tcon cooling boxes with −80°C cooling packs and a dry shipper VOYAGEUR containing absorbed liquid nitrogen, ensuring the cold chain was maintained throughout the journey.

### Ethical considerations and safety measures

The study followed ethical guidelines, with permits from the Department of Wildlife Conservation, Sri Lanka (permit No. WL/3/2/05/18), and ethical clearance from the Institute of Biology, Sri Lanka (ERC IOBSL 170 01 18). Bat handling was conducted following the guidelines of the American Society of Mammalogists, ensuring minimal stress to the animals. Researchers received Pre-exposure rabies prophylaxis (PrEP) and wore personal protective equipment during sample collection to minimize zoonotic risks.

The laboratory analyses of the collected samples were conducted following the BSL-2 conditions at the Robert Koch Institute, Berlin, Germany, adhering to strict safety protocols to minimize any potential risks associated with handling biological materials.

### Laboratory testing

Each bat urine sample was prepared for DNA extraction using a QIAamp viral RNA mini kit (Qiagen, Hilden, Germany), excluding any DNA digestion procedures, in accordance with the manufacturer’s recommendations. Prior to pathogen screening, bat species inhabiting Wavulgalge cave were identified through cytochrome b gene sequence analysis to ensure accurate host species identification.

The extracted DNA was then subjected to PCR amplification with a specific assay targeting the *lipL*32 gene, followed by amplification of positive samples using a reverse primer to obtain longer sequences. Further sequencing of *flaB, secY*, and *rrs2* genes was conducted on positive samples. Purified PCR products were sequenced using the Sanger sequencing method. Details of the assays including primer sequences, PCR conditions, and sequencing protocol are provided in S1 Table.

### Phylogenetic analysis of the sequences

Sequences were analyzed using Geneious Prime® 2023.1.2 and MEGA11 software (*25*). A phylogenetic tree was constructed using MrBayes, with sequences compared to human and bat-associated *Leptospira* species from the NCBI GenBank database.

## Results

Bat species were identified through molecular analysis as *Hipposideros speoris*, *Miniopterus fuliginosus*, *Rhinolophus rouxii*, and *Rousettus leschenaultii* (*26*). A total of 148 urine samples collected from the above four bat species occupying the Wavulgalge cave, Sri Lanka. Among the samples tested, 16 (9.5%) yielded positive results, indicating the presence of *Leptospira* in the Wavulgalge bat population.

The PCR positive samples were identified from *M. fuliginosus* (14/16), *H. speoris* (1/16), and *R. leschenaultii* (1/16) bat species. Summary of the PCR screening results is shown in Table 1.

**Table 1.** Summary of *Leptospira* screening results in four bat species across three sampling periods in Wavulgalge cave, Sri Lanka (March 2018 - January 2019)

| Bat species | 2018 March | 2018 July | 2019 January | Total |
|---|---|---|---|---|
| *H. speoris* | 0/2 | 0/0 | 1/6 | 1/8 |
| *M. fuliginosus* | 0/0 | 14/106 | 0/11 | 14/117 |
| *R. rouxii* | 0/7 | 0/0 | 0/6 | 0/13 |
| *R. leschenaultii* | 0/2 | 0/2 | 1/6 | 1/10 |
| Total | 0/11 | 14/108 | 2/29 | 16/148 |
| Positivity (%) * | 0 | 12.9 | 6.9 | 10.8 |

Bat species, total sample numbers, PCR positive samples as a fraction of the total samples tested per each field visit and per each bat species are shown in the table. \*Due to the low sample numbers, the prevalence rates could not be statistically proven.

### PCR screening results

According to the results, *M. fuliginosus* was tested positive for pathogenic *Leptospir*a bacteria with a 11.97% positivity rate. Positive rates for *H. speoris,* and *R. leschenaultii* could not be calculated due to the insufficient sample numbers. *R. rouxii* urine samples collected during March 2018 and January 2019 were tested negative.

All 11 urine samples collected from *H. speoris, R. leschenaultii* and *R. rouxii* bats during March 2018 were tested negative. The low bat population caused by seasonal migration led to a limited number of bats sampled.

In July 2018, all positive samples were attributed to the highest number of *Miniopterus* bats captured during that field visit. *H. speoris* and *R. rouxii* were not sampled during the July field visit, and the two *R. leschenaultii* bats tested negative.

All four bat species were captured during the January 2019 sampling session. Six urine samples were collected from each bat species, *H. speoris, R. leschenaultii,* and *R. rouxii,* and eleven urine samples were collected from *M. fuliginosus*. However, only one sample from *H. speoris* and one sample from *R. leschenaultii* bats tested positive during the January 2019 sampling session. Both *R. rouxii* and *M. fuliginosus* urine samples tested negative during that session. Nevertheless, statistical inferences could not be drawn from the above data due to the low number of samples from each bat species.

The focus of the study was to conduct a pilot study to explore the potential presence of *Leptospira* in different bat species at specific time points. Thus, drawing definitive conclusions from the data may not be appropriate. Further studies targeting specific questions should be considered for a more comprehensive understanding.

Out of the 16 *Leptospira* PCR positive samples, sequences could be obtained from only 12 samples due to the low DNA amount in 4 samples. Therefore, one positive sample from *R. leschenaultii* and 11 samples from *M. fuliginosus* were sequenced. The resultant sequences have been deposited in the GenBank database at the National Center for Biotechnology Information (NCBI) with the accession numbers given below in Table 2.

**Table 2.**
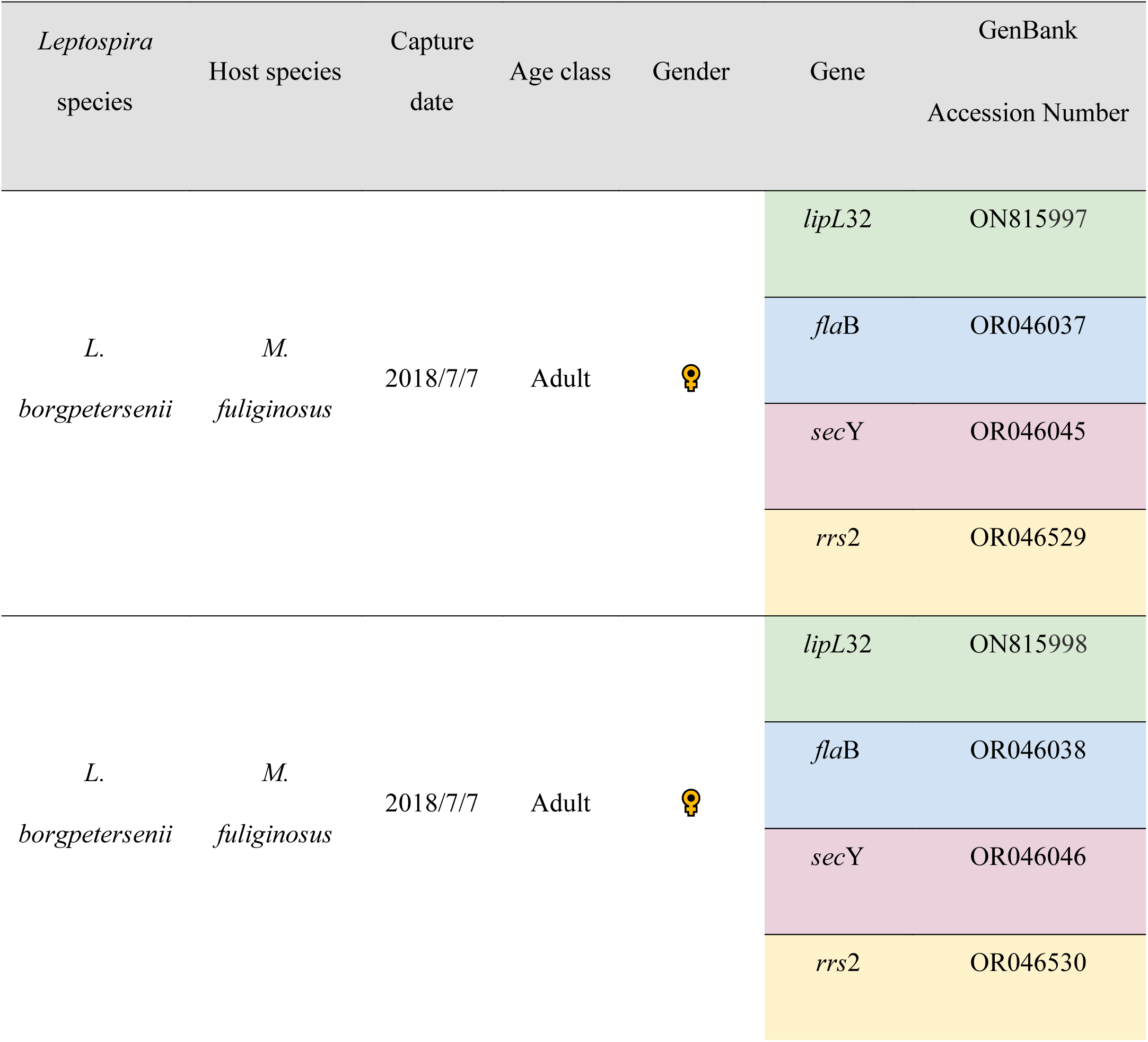
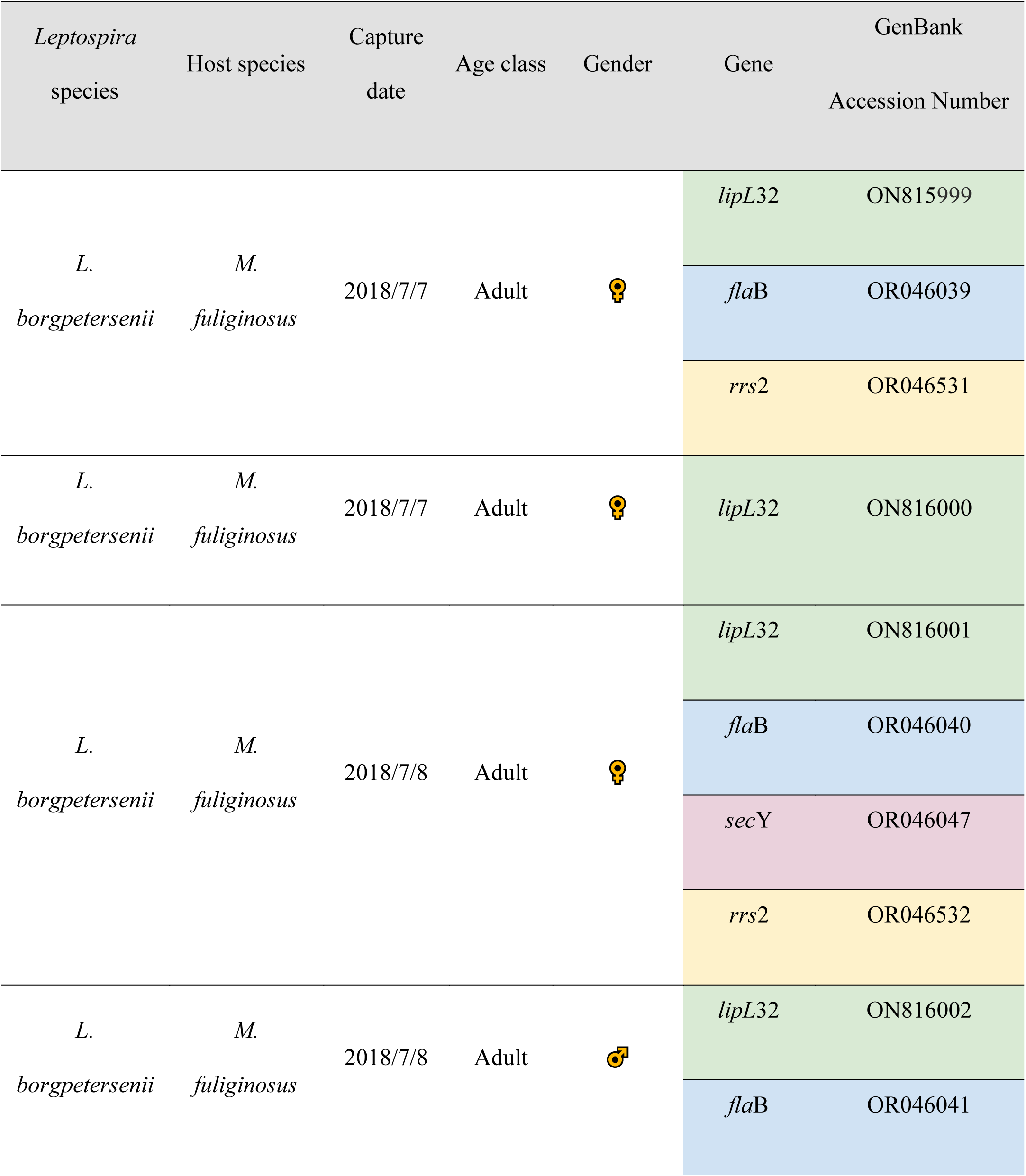
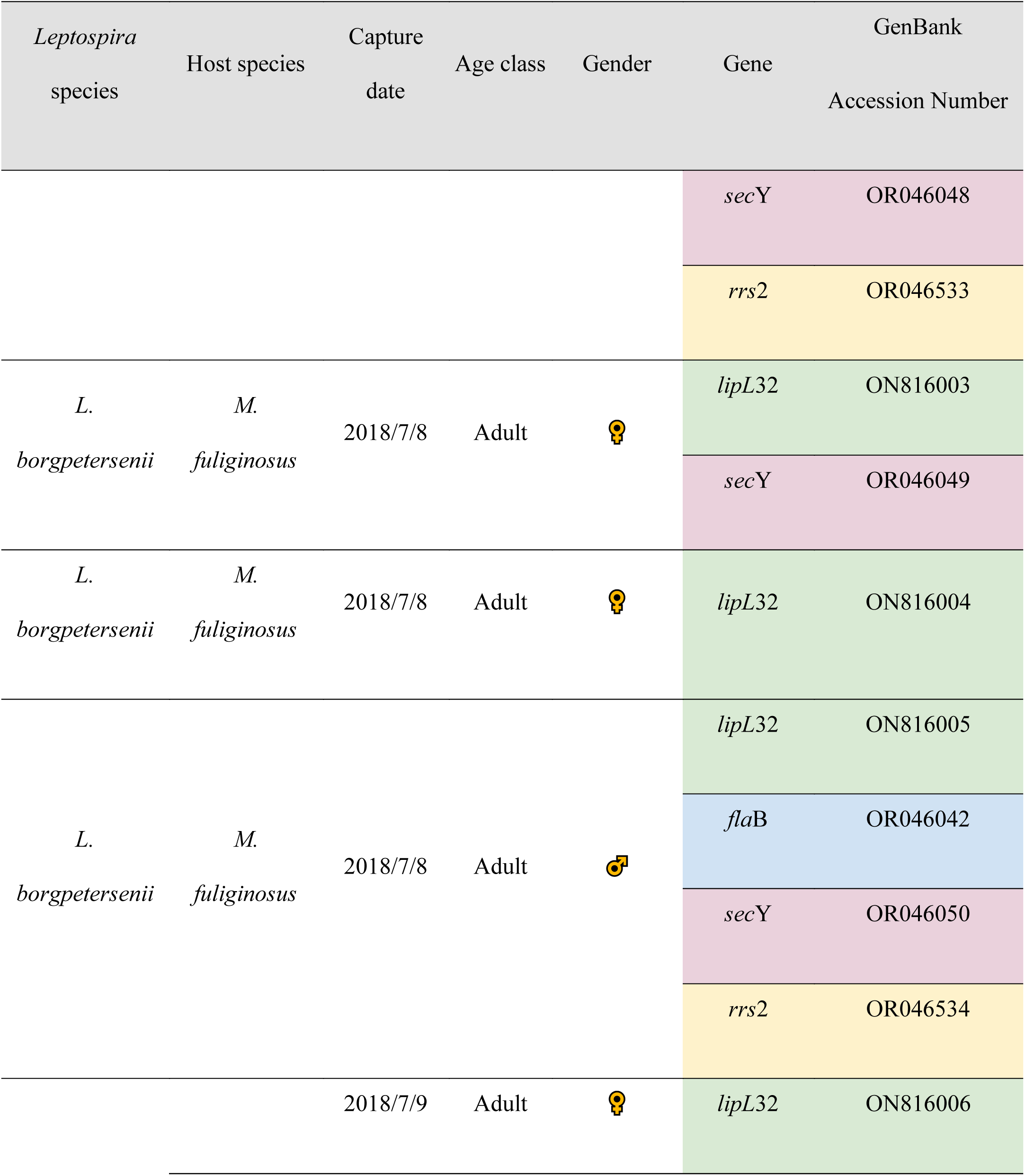
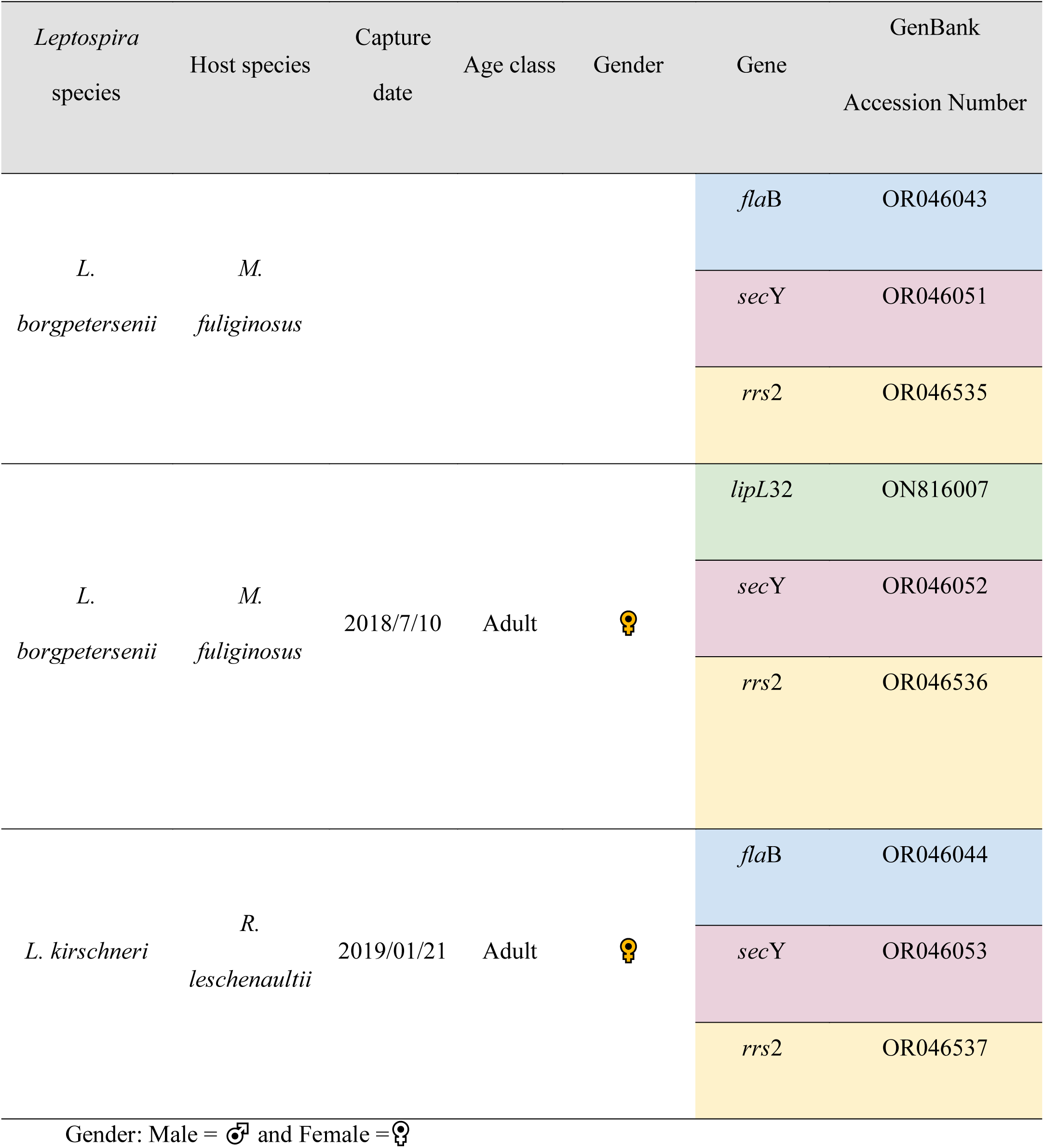
*Leptospira* sequences and associated data for four genes from host bat species captured in Sri Lanka.

| *Leptospira* species | Host species | Capture date | Age class | Gender | Gene | GenBank Accession Number |
|---|---|---|---|---|---|---|
| *L. borgpetersenii* | *M. fuliginosus* | 2018/7/7 | Adult | ♀ | *lipL*32 | ON815997 |
|  |  |  |  |  | *fla*B | OR046037 |
|  |  |  |  |  | *sec*Y | OR046045 |
|  |  |  |  |  | *rrs*2 | OR046529 |
| *L. borgpetersenii* | *M. fuliginosus* | 2018/7/7 | Adult | ♀ | *lipL*32 | ON815998 |
|  |  |  |  |  | *fla*B | OR046038 |
|  |  |  |  |  | *sec*Y | OR046046 |
|  |  |  |  |  | *rrs*2 | OR046530 |
| *Leptospira* species | Host species | Capture date | Age class | Gender | Gene | GenBank Accession Number |
| *L. borgpetersenii* | *M. fuliginosus* | 2018/7/7 | Adult | ♀ | *lipL*32 | ON815999 |
|  |  |  |  |  | *fla*B | OR046039 |
|  |  |  |  |  | *rrs*2 | OR046531 |
| *L. borgpetersenii* | *M. fuliginosus* | 2018/7/7 | Adult | ♀ | *lipL*32 | ON816000 |
| *L. borgpetersenii* | *M. fuliginosus* | 2018/7/8 | Adult | ♀ | *lipL*32 | ON816001 |
|  |  |  |  |  | *fla*B | OR046040 |
|  |  |  |  |  | *sec*Y | OR046047 |
|  |  |  |  |  | *rrs*2 | OR046532 |
| *L. borgpetersenii* | *M. fuliginosus* | 2018/7/8 | Adult | ♂ | *lipL*32 | ON816002 |
|  |  |  |  |  | *fla*B | OR046041 |
| *Leptospira* species | Host species | Capture date | Age class | Gender | Gene | GenBank Accession Number |
|  |  |  |  |  | *sec*Y | OR046048 |
|  |  |  |  |  | *rrs*2 | OR046533 |
| *L. borgpetersenii* | *M. fuliginosus* | 2018/7/8 | Adult | ♀ | *lipL*32 | ON816003 |
|  |  |  |  |  | *sec*Y | OR046049 |
| *L. borgpetersenii* | *M. fuliginosus* | 2018/7/8 | Adult | ♀ | *lipL*32 | ON816004 |
| *L. borgpetersenii* | *M. fuliginosus* | 2018/7/8 | Adult | ♂ | *lipL*32 | ON816005 |
|  |  |  |  |  | *fla*B | OR046042 |
|  |  |  |  |  | *sec*Y | OR046050 |
|  |  |  |  |  | *rrs*2 | OR046534 |
|  |  | 2018/7/9 | Adult | ♀ | *lipL*32 | ON816006 |
| *Leptospira* species | Host species | Capture date | Age class | Gender | Gene | GenBank Accession Number |
| *L. borgpetersenii* | *M. fuliginosus* |  |  |  | *fla*B | OR046043 |
|  |  |  |  |  | *sec*Y | OR046051 |
|  |  |  |  |  | *rrs*2 | OR046535 |
| *L. borgpetersenii* | *M. fuliginosus* | 2018/7/10 | Adult | ♀ | *lipL*32 | ON816007 |
|  |  |  |  |  | *sec*Y | OR046052 |
|  |  |  |  |  | *rrs*2 | OR046536 |
| *L. kirschneri* | *R. leschenaultii* | 2019/01/21 | Adult | ♀ | *fla*B | OR046044 |
|  |  |  |  |  | *sec*Y | OR046053 |
|  |  |  |  |  | *rrs*2 | OR046537 |

Gender: Male = ♂ and Female =♀

### Diversity of *Leptospira* strains among Wavulgalge bat population

All *Leptospira* sequences obtained from *M. fuliginosus* bats were identified as *L. borgpetersenii* species. When compared with the sequences at GenBank database, the *Leptospira* species identified from *M. fuliginosus* from Wavulgalge cave, Sri Lanka showed highest nucleotide (nt) identity for the *Leptospira* species identified from *Miniopterus* bats from Madagascar. The *Leptospira* sequence from *R. leschenaultii* bat was identified as *L. kirschneri* and showed a 95.05% nt identity for *L. kirschneri* identified from a Reed vole from Russia and had a 99.23% nt identity to *Leptospira* species identified from the *Rousettus* bats from Madagascar. BLAST results of all the four genes for all sequences from the study are summarized in the S2 Table.

Phylogenetic trees were constructed for each of the four gene sequences *lipL32*, *flaB*, *secY*, and *rrs2*. The analysis was conducted using the Bayesian algorithm, with 1 million generations, a subsampling frequency of 200, and a burn-in of 10%. The general time reversible (GTR) substitution model was selected with a gamma-distributed rate variation to account for differences in mutation rates across sites. *Leptonema illini* was included as an outgroup to root the trees. Fig 3A, 3B, 3C and 3D show the *lipL32, fla*B, *sec*Y and *rrs*2 gene sequence analyses respectively.

**Fig 3A.**
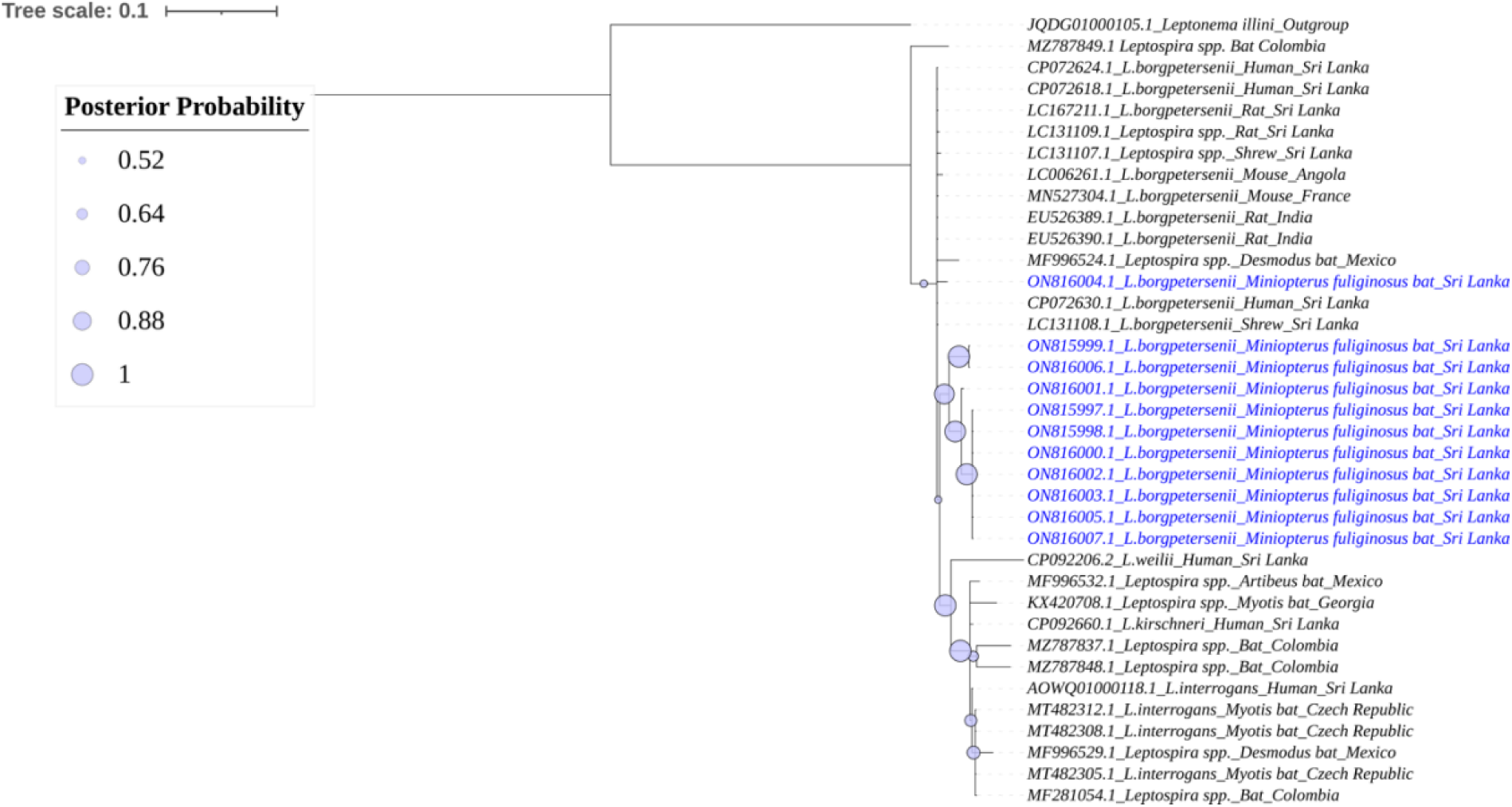
Phylogenetic tree from *lipL32* gene sequences

**Fig 3B.**
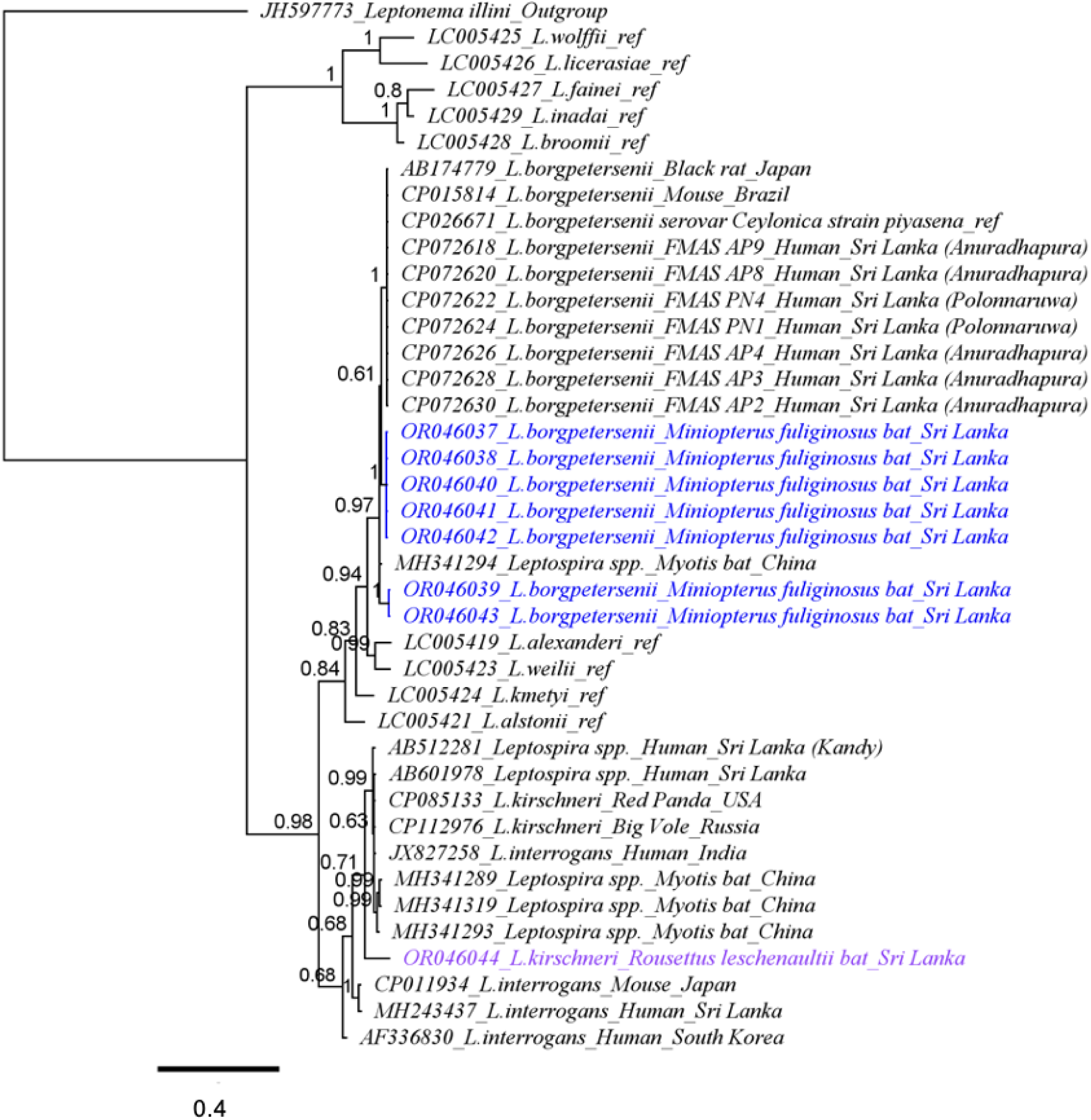
Phylogenetic tree from *fla*B gene sequences

**Fig 3C.**
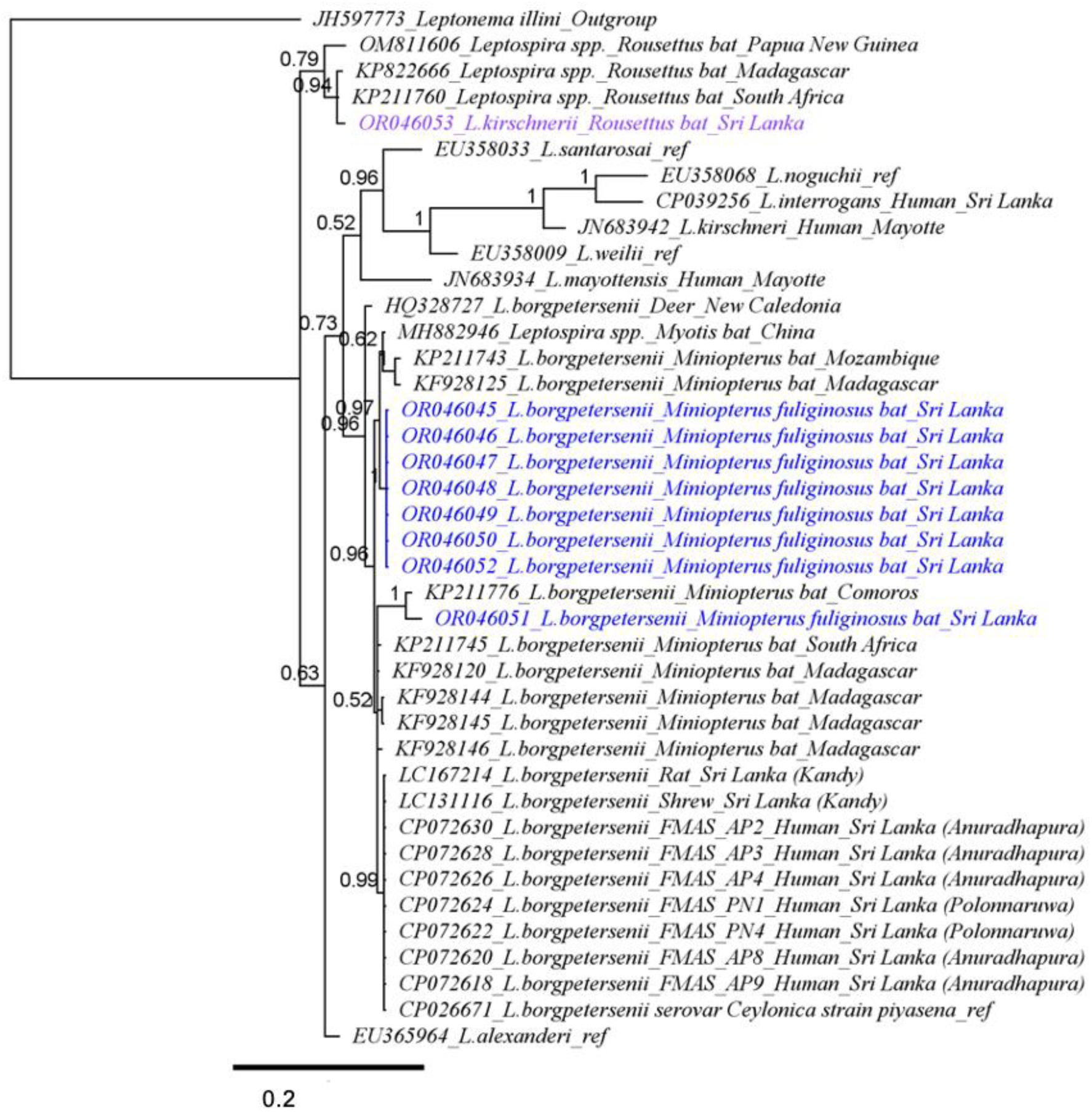
Phylogenetic tree from *secY* gene sequences

**Fig 3D.**
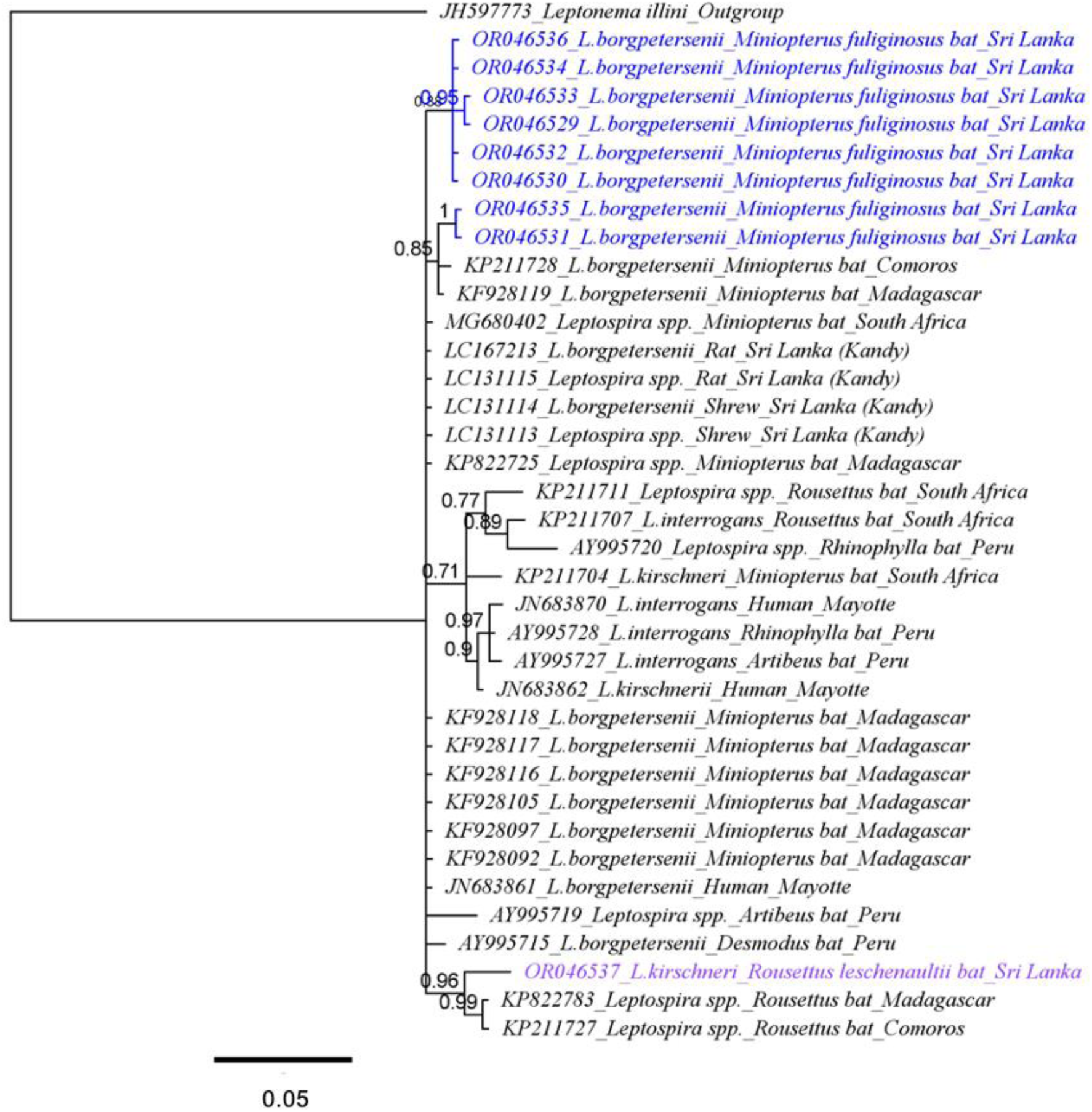
Phylogenetic tree from *rrs2* gene sequences

The phylogenetic tree for the *lipL32* gene, presented in Fig 3A, revealed a close evolutionary relationship between *L. borgpetersenii* strains identified in *Miniopterus* bats from this study to the strain identified from a human from the Anuradhapura district, as well as in shrews from Kandy, Sri Lanka. The human samples were collected between June 2016 and January 2019, while the shrew samples date back to 2015, suggesting a possible transmission link between bats, small mammals, and humans within these regions.

For the *flaB* gene sequences shown in Fig 3B, the *Leptospira* bacteria isolated from *M. fuliginosus* bats exhibited a 96.65% nucleotide identity to *L. borgpetersenii* found in human samples from the Anuradhapura and Polonnaruwa districts. Furthermore, *Leptospira* from *R. leschenaultii* bats showed a 94.92% nucleotide identity to a strain detected in humans from Kandy. These high sequence similarities reinforce the potential zoonotic transmission of *Leptospira* from bats to humans in Sri Lanka.

The phylogenetic analysis of *secY* and *rrs2* gene sequences shown in Fig 3C and Fig 3D respectively, further supports the close evolutionary relationship between *Leptospira* strains isolated from Sri Lankan bats and those found in other regions. The *Leptospira* detected in *Miniopterus* bats were closely related to *L. borgpetersenii* strains isolated from bats of the same genus in South Africa and Madagascar, highlighting the potential global distribution of these *Leptospira* strains within the *Miniopterus* genus. Similarly, *L. kirschneri* bacteria identified in *Rousettus* bats from Sri Lanka showed a strong genetic relationship to strains isolated from *Rousettus* bats in Madagascar, further suggesting cross-regional similarities in bat-borne *Leptospira* species.

Despite the cohabitation of *Miniopterus* and *Rousettus* bats within the Wavulgalge cave, the phylogenetic trees indicate that the *Leptospira* species found in these bats were genetically distinct. This observation points to a potential host-specific adaptation of *Leptospira* strains, as the bats shared the same ecological habitat but harbored different bacterial species. Such host specificity suggests that *Leptospira* strains may be evolving to adapt to particular bat hosts, potentially limiting interspecies transmission within sympatric bat populations.

Overall, these phylogenetic analyses provide important insights into the genetic diversity and evolutionary relationships of *Leptospira* species in Sri Lankan bats, while also highlighting the possible zoonotic and cross-regional transmission of these pathogens. The genetic clustering of *Leptospira* species across different host species and geographic regions underscores the need for further studies to understand the ecology and transmission dynamics of *Leptospira* in bat populations.

## Discussion

This study sheds light on the potential of bats acting as a reservoir for pathogenic *Leptospira* species in Sri Lanka, filling a crucial gap in our understanding of sylvatic cycle of leptospirosis disease transmission in the country. Moreover, the presence of *Leptospira* in bats is of clinical significance, especially in Sri Lanka, where co-infections with Dengue fever are prevalent, particularly during the dengue epidemic season (*27*).

The use of a specific PCR assay targeting the *lipL32* gene, combined with newly designed sequencing primers, was an efficient diagnostic method for species determination of *Leptospira* bacteria among Sri Lankan bats. This study marks the first evidence of *Leptospira* infection within the Wavulgalge bat population, reinforcing the hypothesis that bats could act as important carriers and reservoirs of *Leptospira* in Sri Lanka. These findings highlights the necessity for further research into the sylvatic cycle and transmission dynamics of *Leptospira* to reduce the risk of leptospirosis transmission from bats to humans.

Previous studies have also documented the presence of *Leptospira* in bat species using both immunological and molecular methods (*28–31*). Notably, one case from the USA reported a human contracting leptospirosis after contact with a dead bat in a swimming pool, suggesting a potential risk associated with bat exposure (*32*). This example highlights how direct or indirect interactions with bats can pose significant infection risks.

The host specificity of *Leptospira* bacteria across different regions and bat species is well established (*33*). Research in Madagascar, Mozambique, and South Africa has demonstrated the close association of *L. borgpetersenii* with *Miniopterus* bats, emphasizing the species-specific adaptation of *Leptospira* within certain bat populations (*6,34*). Our study supports these findings, with *L. borgpetersenii* detected in *M. fuliginosus* bats from Sri Lanka, suggesting similar host-specificity patterns.

Conversely, a study focused on *L. borgpetersenii* serogroup *Javanica*, infecting rats (*Rattus spp.*) in Southern Japan, the Philippines, and Taiwan, showed geographical clustering rather than host specificity indicating a broad host range (*35*).

Interestingly, despite the diversity of *Leptospira* species found in the region such as *L. interrogans*, *L. kirschneri*, and *L. weilii*, results from this study showed that all 11 *M. fuliginosus* bats sampled in 2018 were exclusively infected with *L. borgpetersenii* (*36*). Additionally, *L. kirschneri* was detected in a single *R. leschenaultii* bat, closely related to *Leptospira* strains identified in *Rousettus* bats from Madagascar. However, due to the limited sample size, the full extent of host specificity of *Leptospira* in Sri Lankan bats remains unclear.

Furthermore, the study site, Wavulgalge cave, is considered a pre-maternity roost for *Miniopterus* bats, a specific location where pregnant female bats from nearby caves gather before migrating to the maternity roost to give birth (*37*). This increases the likelihood of detecting *Leptospira* in the bat population during the breeding season when large congregations of pregnant females are present. This heightened concentration of bats not only facilitates pathogen transmission within the roost but also poses a risk of spreading *Leptospira* to nearby caves as bats migrate to and from the Wavulgalge cave.

Previous studies indicate two leptospirosis peaks in Sri Lanka, aligning with the two monsoon seasons: a minor peak from March to May and a more significant peak from October to December (*4,38*). The highest number of positive samples was recorded in July, coinciding with the southwestern monsoon and the largest sample collection across the three sampling time points. However, due to the limited sample size in this study, we cannot draw definitive conclusions about the seasonal patterns of *Leptospira* bacterial shedding. In addition, *Leptospira* bacteria can persist in the kidneys of infected bats and be shed intermittently, which may not always coincide with peak transmission periods.

Further research, including longitudinal studies, is needed to better understand the dynamics of *Leptospira* in bats and how it relates to the recognized patterns of leptospirosis outbreaks in Sri Lanka. The specific factors contributing to this observed pattern, such as host immunity, environmental conditions, and reservoir host dynamics, would require more investigation.

Furthermore, the incidence of human leptospirosis cases in Sri Lanka has shown notable variations over the years from 2018 to 2023 as shown in Fig 4 (*39*).

**Fig 4.**
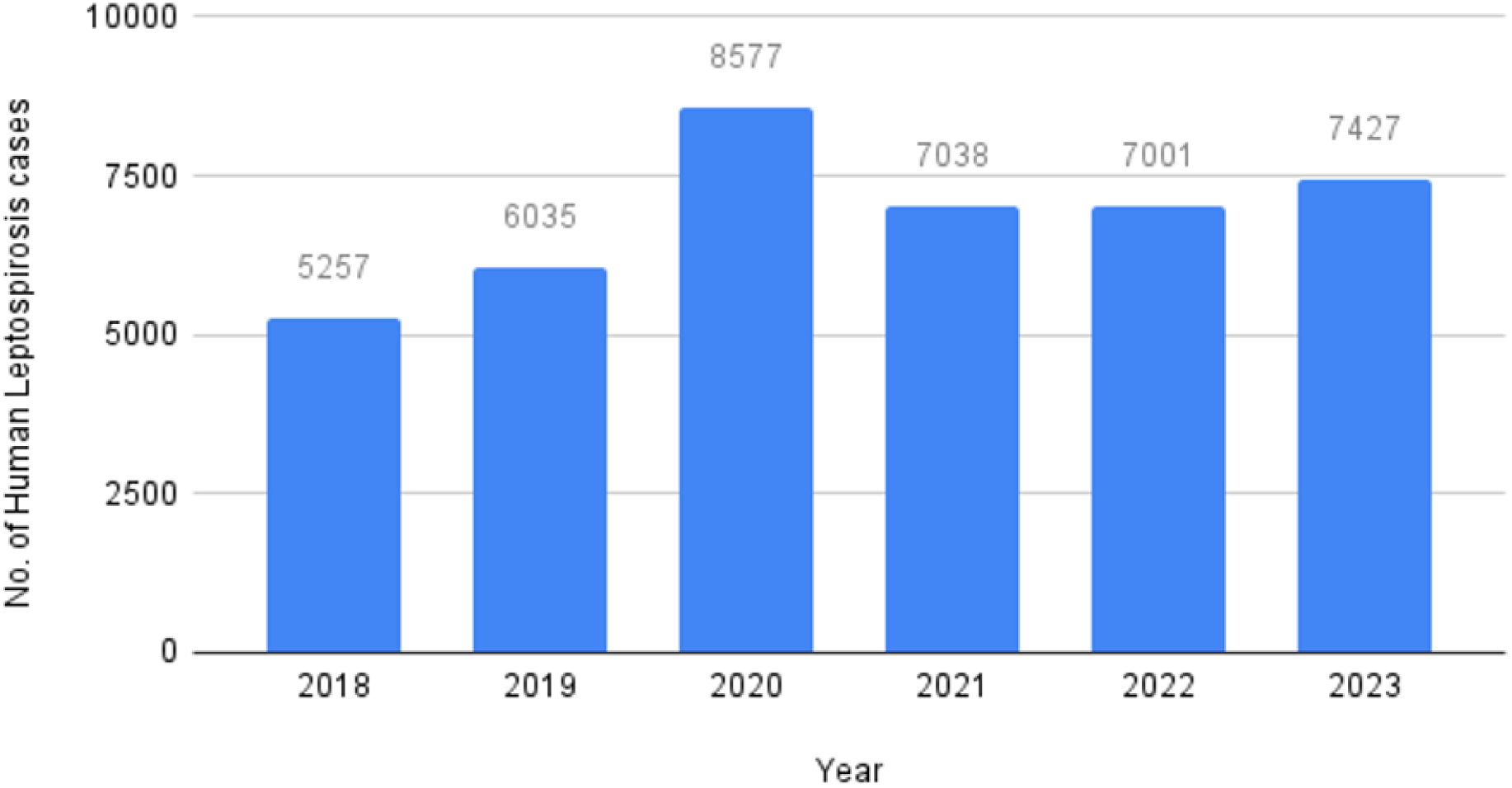
Human leptospirosis cases reported in Sri Lanka from 2018 to 2023. Data source: Epidemiology Unit, Ministry of Health, Sri Lanka.

In 2018, there were 5257 reported cases, which increased to 6035 in 2019. The year 2020 witnessed a significant rise with 8577 cases, marking a substantial surge. However, the trend seemed to slightly decline in 2021, recording 7038 cases, and maintained a similar pattern in 2022 with 7001 reported cases. The data for 2023 indicates a slight increase, reaching 7427 cases. These numbers underscore the dynamic nature of leptospirosis incidence in the country over this period, reflecting potential shifts in environmental and epidemiological factors influencing the prevalence of the disease. Understanding these trends will be crucial for public health initiatives and targeted interventions to mitigate the impact of leptospirosis on the human population.

The correlation observed between the detection of *L. borgpetersenii* DNA in bats and human leptospirosis cases in Sri Lanka is particularly noteworthy, emphasizing the interconnected nature of the *Leptospira* transmission cycle as shown in Fig 5.

**Figure 5.**
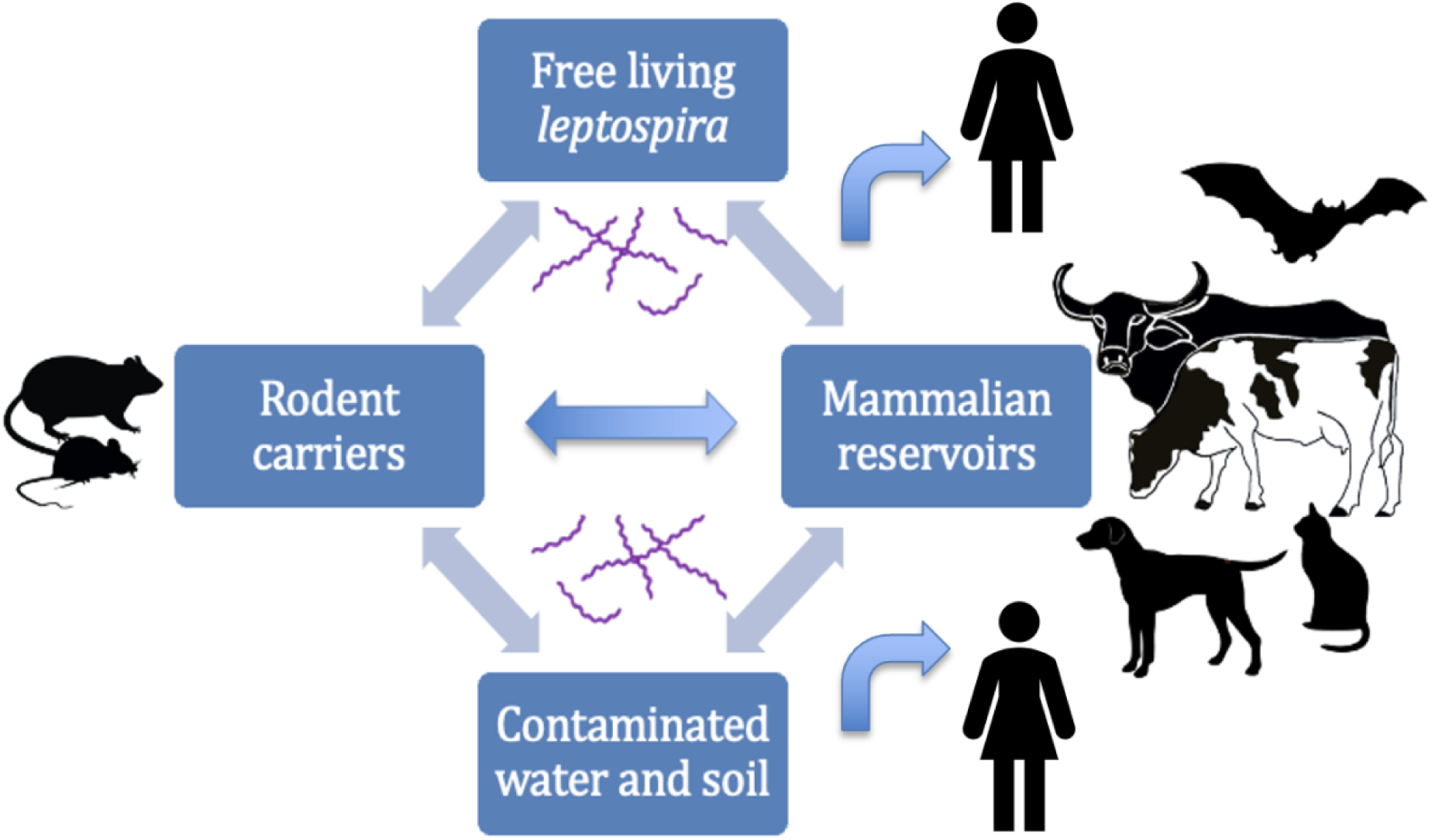
*Leptospira* transmission cycle

Numerous studies suggest that *Leptospira* can persist in bats within moist environments, contaminating other hosts through bat urine (*40,41*). This raises the possibility of *Leptospira* transmission through the use of bat guano as an agricultural fertilizer (*42*). However, the specific tenacity of *Leptospira* in urine or guano may vary depending on factors such as temperature, pH, humidity, exposure to sunlight, and the environmental conditions specific to the location.

Considering the possible transmission routes in Sri Lankan bats, people frequently come into direct contact with bats while collecting guano from the Wavulgalge cave to use as an organic fertilizer in farming. Simultaneously, nearby water bodies are at risk of environmental contamination through bat urination, either directly through dripping or via surface runoff during rain events. Thus, bats could serve as a potential source of direct transmission of *Leptospira* to humans and play a role in the transmission cycle of leptospirosis in the Wellawaya area in Sri Lanka. These factors highlight the need for a One Health approach, integrating human, animal, and environmental health surveillance to control leptospirosis transmission.

Future research is needed in the areas of *Leptospira* transmission mechanisms between bats and humans, bat host specificity, and seasonal variations in prevalence to enhance our understanding of leptospirosis epidemiology. This knowledge will assist public health authorities in designing targeted interventions to mitigate the risk of leptospirosis transmission in Sri Lankan and support the efforts for disease eradication or control.

The strong correlation between the detection of *L. borgpetersenii* DNA in bats and human leptospirosis cases in Sri Lanka is also noteworthy, illustrating the interconnected nature of the *Leptospira* transmission cycle. Although humans are not the primary reservoir of *Leptospira*, they can contract the disease through exposure to contaminated environments or contact with infected animals.

In conclusion, this study not only lays the foundation for our understanding of *Leptospira* dynamics in Sri Lankan bats but also offers insights into the global implications of bats as potential carriers of this zoonotic disease. These findings suggest that Sri Lankan bats play a crucial role as reservoir hosts, maintaining the sylvatic cycle of *Leptospira* bacteria throughout the year and might play a role in occasional outbreaks of the disease.

## Supporting information

S1 Table. Primer sequences and thermocycling conditions for detection of the *lipL32* gene in *Leptospira*

S2 Table. BLAST results of the *Leptospira* sequences obtained from the study

## Author Contributions

Conceptualization, T.P., A.N., W.Y., C.K., S.P. and G.P.; field work and sample collection, T.P., T.M., B.B.Z., S.S. and W.Y.; performing experiments: T.P., and F.S.; formal analysis and interpretation of research data, T.P., F.S., J.W., A.N., W.Y. and C.K.; investigation, T.P., and F.S.; writing—original draft preparation, T.P., C.K., A.N; writing—T.P., F.S., T.M., S.S., B.B.-Z., I.C., S.H., J.W., G.P., S.P., A.N., W.Y. and C.K. review and editing, A.N. and C.K.; visualization, T.P. and C.K.; supervision, S.P., J.W., S.H., F.S., G.P., A.N., W.Y. and C.K.; project administration, A.N. and C.K.; funding acquisition, A.N. and C.K. All authors have read and agreed to the published version of the manuscript.

## Funding

This research was funded by the Federal Ministry of Health, Germany (Bundesministerium für Gesundheit, BMG) under the IDEA (IDentification of Emerging Agents) project within the Global Health Protection Programme (GHPP), grant number ZMVI1-2517-GHP-703 (TP01).

## Institutional Review Board Statement

The animal study protocol was approved by the Department of Wildlife Conservation, Sri Lanka (permit No. WL/3/2/05/18) and the Ethics Committee of the Institute of Biology, Sri Lanka (ERC IOBSL 170 01 18). Animal capturing, handling, and sampling were done in accordance with the Fauna and Flora Protection Ordinance (FFPO) of Sri Lanka.

## Informed Consent Statement

Not applicable.

## Data Availability Statement

The data presented in this study is openly available in GenBank (https://www.ncbi.nlm.nih.gov/genbank).

## Acknowledgments

We thank Angelina Kus, Nicole Kromarek, and Marica Grossegesse for technical and field assistance, and the RKI sequencing lab for providing us with Sanger sequencing results and Timothy Hornsby for the photographs. We convey our gratitude to the Department of Wildlife Conservation, Sri Lanka, and the Institute of Biology, Sri Lanka, for granting us the necessary permits.

## Conflicts of Interest

The authors declare no conflict of interest.

## Notes

### Competing Interest Statement

The authors have declared no competing interest.

## References

1. Terpstra WJ, World Health Organization, International Leptospirosis Society, editors. Human leptospirosis: guidance for diagnosis, surveillance and control. Geneva: World Health Organization; 2003. 109 p.

2. Leptospirosis | CDC [Internet]. 2019 [cited 2023 May 24]. Available from: https://www.cdc.gov/leptospirosis/index.html

3. Epidemiology Unit M of H Sri Lanka. Leptospirosis [Internet]. 2024 [cited 2023 May 24]. Available from: https://www.epid.gov.lk/leptospirosis

4. Ehelepola NDB, Ariyaratne K, Dissanayake WP. The correlation between local weather and leptospirosis incidence in Kandy district, Sri Lanka from 2006 to 2015. Glob Health Action [Internet]. 2019 Jan 1 [cited 2023 Oct 30];12(1):1553283. Available from: https://www.tandfonline.com/doi/full/10.1080/16549716.2018.1553283

5. Agampodi SB, Peacock SJ, Thevanesam V, Nugegoda DB, Smythe L, Thaipadungpanit J, et al. Leptospirosis outbreak in Sri Lanka in 2008: lessons for assessing the global burden of disease. Am J Trop Med Hyg. 2011 Sep;85(3):471–8.

6. Dietrich M, Mühldorfer K, Tortosa P, Markotter W. *Leptospira* and Bats: Story of an Emerging Friendship. Goldman WE, editor. PLOS Pathog [Internet]. 2015 Nov 12 [cited 2023 May 10];11(11):e1005176. Available from: https://dx.plos.org/10.1371/journal.ppat.1005176

7. Calisher CH, Childs JE, Field HE, Holmes KV, Schountz T. Bats: Important Reservoir Hosts of Emerging Viruses. Clin Microbiol Rev [Internet]. 2006 Jul [cited 2023 Sep 21];19(3):531–45. Available from: https://journals.asm.org/doi/10.1128/CMR.00017-06

8. Kohl C, Nitsche A, Kurth A. Update on Potentially Zoonotic Viruses of European Bats. Vaccines [Internet]. 2021 Jun 23 [cited 2023 Jun 19];9(7):690. Available from: https://www.mdpi.com/2076-393X/9/7/690

9. Perera HIT, Yapa WB, Perera HKK. Isolation of *Salmonella* species in *Rousettus leschenaultii* fruit bats in Sri Lanka. In Sri Lanka: Institute of Biology, Sri Lanka; 2016. p. 33. Available from: http://repository.kln.ac.lk/handle/123456789/14527

10. Muzeniek T, Perera T, Siriwardana S, Bas D, Bayram F, Öruc M, et al. Comparative virome analysis of individual shedding routes of *Miniopterus fuliginosus* bats inhabiting the Wavul Galge Cave, Sri Lanka [Internet]. Molecular Biology; 2022 Sep [cited 2023 Jun 23]. Available from: http://biorxiv.org/lookup/doi/10.1101/2022.09.21.508883

11. Athapattu T, Fernando R, Abayawansha R, Fernando P, Fuward M, Samarakoon N, et al. Carrier Status of *Leptospira* spp. in Healthy Companion Dogs in Sri Lanka. Vector-Borne Zoonotic Dis [Internet]. 2022 Feb 1 [cited 2022 Jul 29];22(2):93–100. Available from: https://www.liebertpub.com/doi/10.1089/vbz.2021.0065

12. Denipitiya DTH, Chandrasekharan NV, Abeyewickreme W, Hartskeerl RA, Hapugoda MD. Identification of cattle, buffaloes and rodents as reservoir animals of *Leptospira* in the District of Gampaha, Sri Lanka. BMC Res Notes [Internet]. 2017 Dec [cited 2022 Jul 29];10(1):134. Available from: http://bmcresnotes.biomedcentral.com/articles/10.1186/s13104-017-2457-4

13. Gamage CD, Koizumi N, Perera AKC, Muto M, Nwafor-Okoli C, Ranasinghe S, et al. Carrier Status of Leptospirosis Among Cattle in Sri Lanka: A Zoonotic Threat to Public Health. Transbound Emerg Dis [Internet]. 2014 Feb [cited 2022 Sep 14];61(1):91–6. Available from: https://onlinelibrary.wiley.com/doi/10.1111/tbed.12014

14. Nityananda K, Harvey T. Leptospirosis in Ceylon-Epidemiological and Laboratory Studies. Ceylon J Med Sci [Internet]. 1971 Jun;20(1):5–14. Available from: http://archive.cmb.ac.lk:8080/research/handle/70130/3689

15. Sluydts V, Sarathchandra SR, Piscitelli AP, Van Houtte N, Gryseels S, Mayer-Scholl A, et al. Ecology and distribution of *Leptospira spp*., reservoir hosts and environmental interaction in Sri Lanka, with identification of a new strain. Casanovas-Massana A, editor. PLoS Negl Trop Dis [Internet]. 2022 Sep 16 [cited 2023 May 11];16(9):e0010757. Available from: https://dx.plos.org/10.1371/journal.pntd.0010757

16. Sunil-Chandra NP, Fahlman Å, Waidyarathna S, Näslund J, Jayasundara MVML, Wesula LO, et al. Evidence of orthohantavirus and *Leptospira* infections in small mammals in an endemic area of Gampaha district in Sri Lanka. One Health Outlook [Internet]. 2022 Dec 14 [cited 2023 Oct 30];4(1):17. Available from: https://onehealthoutlook.biomedcentral.com/articles/10.1186/s42522-022-00073-y

17. Haake DA, Zückert WR. The Leptospiral Outer Membrane. In: Adler B, editor. Leptospira and Leptospirosis [Internet]. Berlin, Heidelberg: Springer Berlin Heidelberg; 2015 [cited 2023 May 24]. p. 187–221. (Current Topics in Microbiology and Immunology; vol. 387). Available from: https://link.springer.com/10.1007/978-3-662-45059-8_8

18. Stoddard RA, Gee JE, Wilkins PP, McCaustland K, Hoffmaster AR. Detection of pathogenic *Leptospira spp*. through TaqMan polymerase chain reaction targeting the *LipL32* gene. Diagn Microbiol Infect Dis [Internet]. 2009 Jul [cited 2023 May 22];64(3):247–55. Available from: https://linkinghub.elsevier.com/retrieve/pii/S0732889309001059

19. Ahmed N, Devi SM, De Los Á Valverde M, Vijayachari P, Machang’u RS, Ellis WA, et al. Multilocus sequence typing method for identification and genotypic classification of pathogenic *Leptospira* species. Ann Clin Microbiol Antimicrob [Internet]. 2006 [cited 2023 May 16];5(1):28. Available from: http://ann-clinmicrob.biomedcentral.com/articles/10.1186/1476-0711-5-28

20. Ballados-González GG, Sánchez-Montes S, Romero-Salas D, Colunga Salas P, Gutiérrez-Molina R, León-Paniagua L, et al. Detection of pathogenic *Leptospira* species associated with phyllostomid bats (Mammalia: Chiroptera) from Veracruz, Mexico. Transbound Emerg Dis [Internet]. 2018 Jun [cited 2022 Jul 29];65(3):773–81. Available from: https://onlinelibrary.wiley.com/doi/10.1111/tbed.12802

21. Kamani J, Harrus S, Ocholi RA, Yague II, Nyango PG, González-Miguel J, et al. Molecular detection and characterization of pathogenic *Leptospira* species in bats (Chiroptera) roosting in human habitats in Nigeria, West Africa. Zoonoses Public Health [Internet]. 2021 Dec [cited 2022 Jul 29];68(8):908–16. Available from: https://onlinelibrary.wiley.com/doi/10.1111/zph.12880

22. Zhao M, Xiao X, Han HJ, Wang LJ, Lei SC, Liu JW, et al. *Leptospira* in Bats from Hubei Province, China, 2018. J Wildl Dis [Internet]. 2019 Oct 9 [cited 2022 Jul 29];55(4):940. Available from: https://bioone.org/journals/journal-of-wildlife-diseases/volume-55/issue-4/2019-01-009/Leptospira-in-Bats-from-Hubei-Province-China-2018/10.7589/2019-01-009.full

23. secY - SecY - *Leptospira interrogans* | UniProtKB | UniProt [Internet]. [cited 2023 May 24]. Available from: https://www.uniprot.org/uniprotkb/A0A089NBG6/entry

24. flaB - Flagellin - *Leptospira borgpetersenii* | UniProtKB | UniProt [Internet]. [cited 2023 May 24]. Available from: https://www.uniprot.org/uniprotkb/Q9ZHF9/entry

25. Tamura K, Stecher G, Kumar S. MEGA11: Molecular Evolutionary Genetics Analysis Version 11. Battistuzzi FU, editor. Mol Biol Evol [Internet]. 2021 Jun 25 [cited 2023 May 30];38(7):3022–7. Available from: https://academic.oup.com/mbe/article/38/7/3022/6248099

26. Perera T, Schwarz F, Muzeniek T, Siriwardana S, Becker-Ziaja B, Perera IC, et al. First Complete Cytochrome B Sequences and Molecular Taxonomy of Bat Species from Sri Lanka. Animals [Internet]. 2022 Jun 29 [cited 2023 May 30];12(13):1674. Available from: https://www.mdpi.com/2076-2615/12/13/1674

27. Samarakoon PSMJU, Karunanayake L, Muthugala R, Karunanayake P. Prevalence, clinical, and demographic characteristics of concomitant dengue fever and leptospirosis among acute dengue fever patients in The Western Province of Sri Lanka. WHO South-East Asia J Public Health. 2022;11(2):93–6.

28. Liceras de Hidalgo J. Leptospirosis in Tingo Maria, Huanuco Department, Peru. II. Study in wild animals. Boletin Oficina Sanit Panam Pan Am Sanit Bur. 1981;91(1):47–55.

29. Bunnell JE, Hice CL, Watts DM, Montrueil V, Tesh RB, Vinetz JM. Detection of pathogenic *Leptospira* spp. infections among mammals captured in the Peruvian Amazon basin region. Am J Trop Med Hyg. 2000;63(5):255–8.

30. Smythe LD, Field HE, Barnett LJ, Smith CS, Dohnt MF, Symonds ML, et al. Leptospiral antibodies in flying foxes in Australia. J Wildl Dis. 2002;38(1):182–6.

31. Matthias MA, Díaz MM, Campos KJ, Calderon M, Willig MR, Pacheco V, et al. Diversity of bat-associated *Leptospira* in the Peruvian Amazon inferred by bayesian phylogenetic analysis of 16S ribosomal DNA sequences. Am J Trop Med Hyg. 2005 Nov;73(5):964–74.

32. Vashi NA, Reddy P, Wayne DB, Sabin B. Bat-Associated Leptospirosis. J Gen Intern Med [Internet]. 2010 Feb [cited 2022 Sep 14];25(2):162–4. Available from: http://link.springer.com/10.1007/s11606-009-1210-7

33. Cordonin C, Turpin M, Bringart M, Bascands JL, Flores O, Dellagi K, et al. Pathogenic *Leptospira* and their animal reservoirs: testing host specificity through experimental infection. Sci Rep [Internet]. 2020 Apr 29 [cited 2023 May 12];10(1):7239. Available from: https://www.nature.com/articles/s41598-020-64172-4

34. Gomard Y, Dietrich M, Wieseke N, Ramasindrazana B, Lagadec E, Goodman SM, et al. Malagasy bats shelter a considerable genetic diversity of pathogenic *Leptospira* suggesting notable host-specificity patterns. Olson J, editor. FEMS Microbiol Ecol [Internet]. 2016 Apr [cited 2022 Jul 29];92(4):fiw037. Available from: https://academic.oup.com/femsec/article-lookup/doi/10.1093/femsec/fiw037

35. Koizumi N, Wada T, Morita M, Mu JJ, Ohnishi M. Comparative genomic analysis of *Leptospira borgpetersenii* serogroup Javanica isolated from *Rattus* species in Southern Japan, Philippines, and Taiwan. Infect Genet Evol [Internet]. 2020 Nov [cited 2022 Jul 29];85:104447. Available from: https://linkinghub.elsevier.com/retrieve/pii/S1567134820302781

36. Jayasundara D, Senavirathna I, Warnasekara J, Gamage C, Siribaddana S, Kularatne SAM, et al. Correction: 12 Novel clonal groups of *Leptospira* infecting humans in multiple contrasting epidemiological contexts in Sri Lanka. PLoS Negl Trop Dis [Internet]. 2021 May 26 [cited 2023 May 12];15(5):e0009471. Available from: https://dx.plos.org/10.1371/journal.pntd.0009471

37. Yapa WB, Digana PMCB, Ratnasooriya WD, Rübsamen R, Costa HH, Randeniya PV. Breeding associated migration of *Miniopterus schreibersii* between two natural caves in Sri Lanka. In: Proceedings of the 19h Annual Sessions of the Institute of Biology, Sri Lanka. Sri Lanka: Institute of Biology,; 1998. p. P21.

38. Robertson C, Nelson TA, Stephen C. Spatial epidemiology of suspected clinical leptospirosis in Sri Lanka. Epidemiol Infect [Internet]. 2012 Apr [cited 2023 Nov 2];140(4):731–43. Available from: https://www.cambridge.org/core/product/identifier/S0950268811001014/type/journal_article

39. Samaraweera DS. Trends of Leptospirosis in Sri Lanka for the year 2022 Part I. 2023;50.

40. Joffrin L, Dietrich M, Mavingui P, Lebarbenchon C. Bat pathogens hit the road: But which one? Dutch RE, editor. PLOS Pathog [Internet]. 2018 Aug 9 [cited 2023 May 12];14(8):e1007134. Available from: https://dx.plos.org/10.1371/journal.ppat.1007134

41. Thibeaux R, Geroult S, Benezech C, Chabaud S, Soupé-Gilbert ME, Girault D, et al. Seeking the environmental source of Leptospirosis reveals durable bacterial viability in river soils. PLoS Negl Trop Dis. 2017 Feb;11(2):e0005414.

42. Newman SH, Food and Agriculture Organization of the United Nations, editors. Investigating the role of bats in emerging zoonoses: balancing ecology, conservation and public health interest. Rome: Food and Agriculture Organization of the United Nations; 2011. 169 p. (FAO animal production and health manual).

